# Efficient gene activation in plants by the MoonTag programmable transcriptional activator

**DOI:** 10.1101/2023.02.15.528671

**Authors:** J. Armando Casas-Mollano, Matthew Zinselmeier, Adam Sychla, Michael J Smanski

## Abstract

CRISPR/Cas-based transcriptional activators have been developed to induce gene expression in eukaryotic and prokaryotic organisms. The main advantages of CRISPR-Cas based systems is that they can achieve high levels of transcriptional activation and are very easy to program via pairing between the guide RNA and the DNA target strand. SunTag is a second-generation system that activates transcription by recruiting multiple copies of an activation domain (AD) to its target promoters. SunTag is a strong activator; however, in some species it is difficult to stably express. To overcome this problem, we designed MoonTag, a new activator that worked on the same basic principle as SunTag, but whose components are better tolerated when stably expressed in transgenic plants. We demonstrate that MoonTag is capable of inducing high levels of transcription in all plants tested. In Setaria, MoonTag is capable of inducing high levels of transcription of reporter genes as well as of endogenous genes. More important, MoonTag components are expressed in transgenic plants to high levels without any deleterious effects. MoonTag is also able to efficiently activate genes in eudicotyledonous species such as Arabidopsis and tomato. Finally, we show that MoonTag activation is functional across a range of temperatures, which is promising for potential field applications.

## Introduction

CRISPR/Cas-based transcriptional activators have been developed to induce gene expression in eukaryotic organisms. CRISPR/Cas-based activators comprise two main components: (i) a DNA binding domain which consists of a catalytically inactive or nuclease “dead” Cas9 (dCas9) protein, and (ii) an activation domain (AD) capable of stimulating transcription when targeted a core promoter region (1). The main advantages of CRISPR/Cas-based systems is that they can achieve high levels of transcriptional activation and are very easy to program via pairing between the guide RNA and the DNA target strand (1). The first-generation CRISPR-Cas activators were created by direct fusion of an AD to the C-terminus of the dCas9 protein. The resulting activator was capable of inducing the expression of reporters and endogenous genes albeit to moderate levels (2–4). A more efficient, second-generation of CRISPR/Cas activators was created by constructing systems that recruit multiple ADs to the dCas9 ribonucleoprotein (1). Several approaches have been used for the recruitment of multiple ADs. For instance, in the dCas9-TV system several copies of the TAL and VP64 ADs are fused in tandem to dCas9 (5). Scaffold RNA (scRNA) and synergistic activation mediator (SAM) uses a sgRNA modified with hairpins that allow the use of RNA-binding proteins to tether ADs to dCas9 ribonucleoprotein (3, 6, 7). In the SunTag system the recruitment of multiple copies of an AD is achieved using an antigen-antibody interaction between a single-chain fragment variable (scFv, fused to the VP64 AD) and a GCN4 peptide (a 19 amino acid peptide fused to the dCas9 protein) (8). SunTag was designed to contain 10 copies of the GCN4 peptide fused to dCas9 being capable to potentially recruit up to 10 copies of the VP64 domain to the target promoter (8). The three-component repurposed technology for enhanced expression (TREE) improves the recruitment of multiple ADs by combining the RNA tethering system of SAM with the SunTag scFv-GCN4 interaction tags (9).

Both first- and second-generation of CRISPR/Cas activators has been tested in several plant species. dCas9-TV was shown to be capable of activating endogenous genes when transiently expressed in Arabidopsis and rice protoplasts (5). Activation of endogenous genes and the expected overexpression phenotypes were observed in Arabidopsis and transgenic plants expressing dCas9-TV (10). Several iterations of SAM, scRNA and SunTag were comparatively tested in a transient system in *Nicotiana benthamiana* in order to identify combinations of ADs producing stronger activation (11). In transgenic plants, endogenous genes targetted by SAM activators could drive sufficient levels of overexpression to induce a phenotype (12, 13). Similarly, SunTag has been tested in Arabidopsis, showing strong transcriptional activation of endogenous genes with the occurrence of the expected phenotypes (14). Recently, a TREE system, ACT 3.0, was shown capable of strong gene activation in protoplast transient systems and in transgenic plants from Arabidopsis, tomato and rice (15).

Given that SunTag performed very well as an activator in Arabidopsis we decided to test its use in monocot plant species. However, out initial attempts to express SunTag in *Setaria viridis* transgenic plants resulted in plants expressing low levels on the scFv component of SunTag (*vide infra*). For this reason we decided to replace the scFv-GCN4 interaction tags for a new pair that will be better expressed in plants but still functions as a strong activator in plants. Inspired by previous studies that demonstrate recruitment of fluorescent proteins via a similar mechanism (16), we develop a new sequence-programmable transcriptional activator, named MoonTag, that uses a nanobody NbGP41 and a GP41 peptide pair to mediate the recruitment of the VP64 activation domain. Here we show that MoonTag is well-tolerated when expressed in transgenic plants where it can efficiently activate gene expression from synthetic promoters and also endogenous genes. We present data from diverse species and target genes that cumulatively show broad utility of MoonTag activators to control gene expression in plants.

## Results

### Design of the MoonTag-VP64 system for gene activation in Setaria

We initially set out to test transcriptional activation in Setaria using the SunTag system. SunTag was shown before to been able to induce high levels of gene activation in Arabidopsis (14). We use the SunTag system previously reported in Arabidopsis but with 10X and 24X copies of the GCN4 peptide (ST1-10X and ST1-24X, respectively) and place the coding sequence of the components under the control of the CmYLCV promoter (17) to drive expression in Setaria (Supplementary Figure S1). An additional SunTag variant (ST2) was created by removing the sfGFP coding sequence of the scFv module (Supplementary Figure S1). We tested the different SunTag versions for transient activation of an inducible promoter driving a luciferase reporter in Setaria protoplasts described previously (18). This assay showed that both versions of SunTag were capable of inducing the expression of the luciferase reporter. However, only low levels of luciferase activity was observed with ST1 whereas the use of ST2 led to high induction of luciferase activity (Figure 1a). Additionally, use of ST2 with 24X copies of GCN4 lead to the highest luciferase activity when compared with the same ST2 with 10X copies of GCN4. Given that ST2-24X was the most efficient version of SunTag, a binary vector containing the dCas9-24XGCN4 and the scFv-VP64-GB1 components of SunTag driven by the cmYLCV promoter was constructed and used to generate transgenic Setaria plants. Setaria transformation with the binary vector expressing the ST2 components resulted in only two transgenic lines obtained after two large transformation attempts. RT-qPCR analysis of plants from these two lines indicated that while the dCas9-24XGCN4 is expressed to moderate levels, the expression of the scFv-VP64-GB1 component is considerable low in both lines (Supplementary Figure S2). We hypothesize that poor expression of the scFv-VP64-GB1 gene is due to transgene silencing that is likely driven by the cellular stress caused by expressing the scFv component, which is notoriously difficult to express (see discussion below). Therefore, we decided to retool SunTag with a new antigen-antibody pair more amenable to expression in plants. Boersma and colleagues recently carried out an screen for antigen-antibody pairs to develop a fluorescent labelling system orthologous to SunTag (16). We used one of these antigen-antibody pair to create a new activator called MoonTag keeping with the naming of the tag used by Boersma and colleagues. The MoonTag activator replaces the scFv-GCN4 antibody-epitope interaction of SunTag with a llama nanobody GP41-GP41 peptide antibody-epitope interaction to recruit the VP64 AD (Figure 1b). NbGP41 (clone 2H10) in MoonTag, is a llama nanobody that recognizes a 15 amino acid peptide (N-KNEQELLELDKWASL-C) from the membrane proximal external region (MPER) of the HIV-1 glycoprotein gp41 (19). The smaller and more soluble nanobody GP41, as a fusion with sfGFP and GB1, seems to be readily expressed in Setaria protoplasts (Supplementary Figure S3). The diagram in Figure 1b shows the components of the MoonTag activating system. It includes (i) a DNA-binding component comprising dCas9 fused to several copies (10 copies in the Figure) of the 15-residue GP41 peptide (dCas9-10XGP41), (ii) an activation module comprising the nanobody GP41 fused to the sfGFP (super folder GFP), the VP64 activation domain and the GB1 solubility tags (NbGP41-sfGFP-VP64-GB1), and (iii) a sgRNA that directs the assembly to a gene of interest. When expressed in plant cells dCas9-10XGP41 is able to bind to its target regions guided by the sgRNA. The GP41 peptide repeats in dCas9-10XGP41 interact with the GP41 nanobody of NbGP41-sfGFP-VP64-GB1, thus recruiting up to 10 copies of the VP64 activation domain to the ribonucleoprotein complex (Figure 1b). To further develop the MoonTag activation system, we tested several design variations and measured target gene activation. Specifically, we tested dCas9 fused to different number (10X or 24X) of repeats of the GP41 peptide (MT1). We also created a version of the activation module lacking sfGFP (NbGP41-VP64-GB1; MT2) and lacking both sfGFP and GB1 (NbGP41-VP64; MT3) solubility tags. MoonTag variants were first tested for their ability to activate a promoter driving luciferase in a Setaria protoplast transient expression system. Activation of the promoter, as determined by luciferase activity, was observed in the presence of all MoonTag constructs tested, albeit to different levels (Figure 1a). Luciferase activity was the highest with MoonTag with the NbGP41-VP64-GB1 component, whereas the lowest activity was observed with MoonTag with NbGP41-sfGFP-VP64-GB1. Increased copy number of the GP41 peptide fused to dCas9 seems to lead to higher activation levels but not in the MoonTag version with NbGP41-VP64-GB1. When comparing MoonTag activation to that of equivalent SunTag constructs, MoonTag seems to provide similar or slightly better activation levels than SunTag (Figure 1a). Thus, these results suggest that MoonTag works as a transcriptional activator capable of induce expression of a reporter gene to similar levels than the SunTag activating system.

**Fig. 1.**
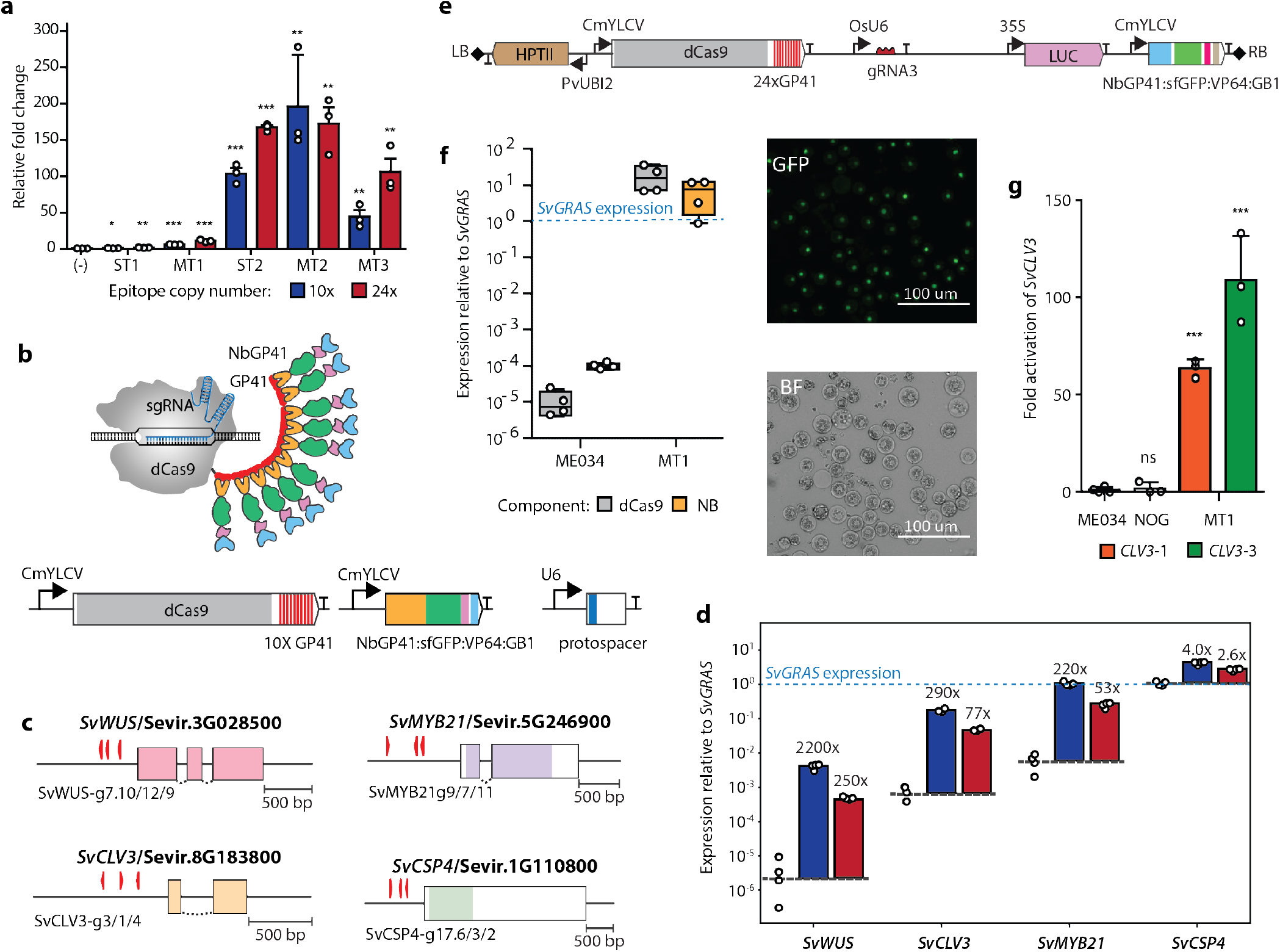
Characterization of the MoonTag activator in Setaria. (a) Activation of a luciferase reporter by SunTag and MoonTag in Setaria protoplasts. Expression of SunTag without the scFv component is used as control. Blue and red bars indicate 10x or 24x copies of the epitope peptides fused to dCas9, respectively. SunTag (ST1-2) and MoonTag (MT1-3) designs are described in the main text. (b) Schematic representation of the MoonTag activator (top) and diagrams of the expression constructs encoding the components of MoonTag. (c) Exon map of genomic targets, *SvWUS, SvMYB21, SvCLV3*, and *SvCSP4* with sgRNA binding sites annotated by right-facing (template strand) or left-facing (complement strand) arrows. Green arrow for *SvCLV3* correspond to panel (g). (d) Activation of the indicated endogenous genes by MoonTag in Setaria protoplasts. No-sgRNA controls (left-most datapoints for each target gene) indicate basal expression levels compared to the reference gene, *SvGRAS*. Blue and red bars represent epitope number, as in (a). Fold-change in expression relative to basal levels is given atop each bar. (e) T-DNA of the binary vector constructed to express the MoonTag activator in Setaria. The genes encoding the different components of MoonTag are indicated. dCas9-24XGP41 and NbGP41-sfGFP-VP64-GB1 are driven by the CmYLCV promoter. The SvCLV3 gRNA, indicated as gRNA3, is driven by the rice U6 promoter. Expression of the selectable marker HPTII is driven by the switchgrass UBI2 promoter. A Luciferase reporter driven by the 35S promoter was also included in the binary vector. Right (RB) and left borders (LB) of the T-DNA are represented by grey rhombi. (f) Left: expression of MoonTag components, dCas9-24×GP41 and NbGP41-sfGFP-VP64-GB1, relative to the reference gene *SvGRAS* in leaves of four putative T0 transgenic plants. Right: representative GFP signal from NbGP41-sfGFP-VP64-GB1 in protoplasts prepared from leaves of transgenic plants expressing the MoonTag. GFP: Fluorescent microscopy. BF: bright field microscopy. (g) Expression of *SVCLV3* in wild type (ME034), no-sgRNA control (NOG), and homozygous T2 Setaria transgenic plants. For each transgenic line, gRNA3-1 and gRNA3-3, the leaves of 3 homozygous individuals were analyzed. *SvCLV3* expression is expressed as average of three individuals for each line and normalized to its expression in the wild type ME034.

### MoonTag-VP64 system is able to activate endogenous genes in Setaria

We next sought to investigate if MoonTag was capable of activating endogenous genes in Setaria. We designed three sgRNAs within 500 bp of the promoter region of four genes, *SvWUSCHEL* (*SvWUS*), *SvCLAVATA3* (*SvCLV3*), *SvMYB21* and *SvCSP4* (Figure 1c). Transient expression of MoonTag in the presence of sgRNAs binding to the promoter of endogenous genes lead to the increased expression of all targets (Figure 1d). Large increases in activation were observed for lowly expressed genes such as *SvWUS*, *SvCLV3* and *SvMYB21* whereas only modest levels of activation were observed for moderately expressed gene *SvCSP4* (Figure 1d). This agrees with previous observations suggesting that genes with high basal expression levels are less efficiently activated by CRISPR/Cas activators (Casas-Mollano et al., 2020). Taken together, these observations indicate that MoonTag is an efficient activator of transgenes and endogenous genes when transiently expressed in Setaria protoplasts. We next investigated the performance of MoonTag activators in stable transgenic Setaria plants. Because Setaria can be transformed using Agrobacterium, we constructed a binary vector capable of expressing MoonTag with a single sgRNA that targets the promoter of the *SvCLV3* gene. This construct contains dCas9-24XGP41 and NbGP41-sfGFP-VP64-GB1 each driven by the CmYLCV promoter, the CLV3-sgRNA driven by a rice U6 polymerase III promoter, and a HPTII selectable marker conferring resistance to hygromycin (Figure 1e). After Agrobacterium-mediated transformation, several hygromycin resistant plants were obtained and transferred to soil. Plants transferred to soil did not show any growth defects or obvious morphological changes. We measured the expression of dCas9-24XGP41 and NbGP41-sfGFP-VP64-GB1 in four of these lines and found that all plants expressed the MoonTag components at levels greater than the *SvGRAS* reference gene (Figure 1f). A GFP signal from the expression of NbGP41-sfGFP-VP64-GB1 was observed in the nuclei of protoplasts produced from leaves of MoonTag-expressing plants (Figure 1f). Two transgenic lines showing the highest levels of *SvCLV3* activation were then grown for two generations to produce homozygous seeds. Expression of *SvCLV3* in homozygous lines was shown to be between 50 to 100 fold higher than in the wild-type ME034 control (Figure 1g). Expression of the MoonTag components (dCas9-24XGP41 and NbGP41-sfGFP-VP64-GB1) and the CLV3-gRNA3 were also maintained in the homozygous lines (Supplementary Figure S4). In spite of the strong *SvCLV3* activation, no obvious growth or developmental phenotypes were observed in plants of the two homozygous lines overexpressing *SvCLV3*, nor in the no-sgRNA MoonTag control line. (Supplementary Figure S5). *SvCLV3* overexpression produces a clear phenotype in dicots (20), but its overexpression in Setaria has not been characterized. Cumulatively, these observations suggest that Setaria transgenic plants stably expressing high levels of MoonTag components can be obtained without deleterious effects. More important, transgenic plants expressing MoonTag targeted to the SvCLV3 promoter by a single sgRNA resulted in the increased expression of this endogenous gene in Setaria.

### MoonTag-driven expression of a luciferase reporter in tomato hairy roots

We next sought to study the ability of MoonTag to activate genes in eudicotyledonous species such as tomato and Arabidopsis. In tomato, we decided to use hairy roots produced by *Agrobacterium rhizogenes* to compare the MoonTag and SunTag activators. For hairy root generation we made constructs that contained a luciferase reporter driven by a synthetic promoter activated by either MoonTag or SunTag (Figure 2a). As a control, a construct expressing all components of MoonTag but without a gRNA (NOG) targeting the promoter of the reporter were generated. After transformation, the hairy roots obtained with the different constructs were analyzed for luciferase activity and the expression of the luciferase reporter and the activator components. Hairy roots treated with D-luciferin showed increased luminescence signal in plants transformed with MoonTag and SunTag compared to that of the control roots expressing MoonTag without a gRNA (Figure 2b). Similarly, expression analysis in hairy roots indicate that both activators express the luciferase reporter to higher levels than roots expressing MoonTag without a gRNA (Figure 2c). However, Luciferase expression in hairy roots transformed with MoonTag was in general more efficient than that obtained with SunTag. One caveat to this observation is that the SunTag version used contains 10 copies of the GCN4 peptide whereas the MoonTag versions used contains 13 and 24 copies of the GP41 peptide. Ten months after continuous subculture, expression of the luciferase is maintained in both SunTag and MoonTag constructs suggesting that gene activation mediated by both activators is stable over many mitotic cell divisions (Figure 2d). Expression analysis also indicated that while the expression of dCas9 is generally lower in SunTag, the expression of the antibody fusion is similar in both activators (Supplementary Figure S6). In contrast, the expression of the gRNAs varies widely in both activators and among different constructs (Supplementary Figure S6). Furthermore, the GFP signal of the antibody components (scFv-sfGFP-VP64-GB1 for SunTag and NbGP41-sfGFP-VP64-GB1 for MoonTag) is usually higher in MoonTag than SunTag (Supplementary Figure S7). This suggest that even though both components are expressed at the same level, the GP41 nanobody can accumulate to higher levels in tomato hairy roots.

**Fig. 2.**
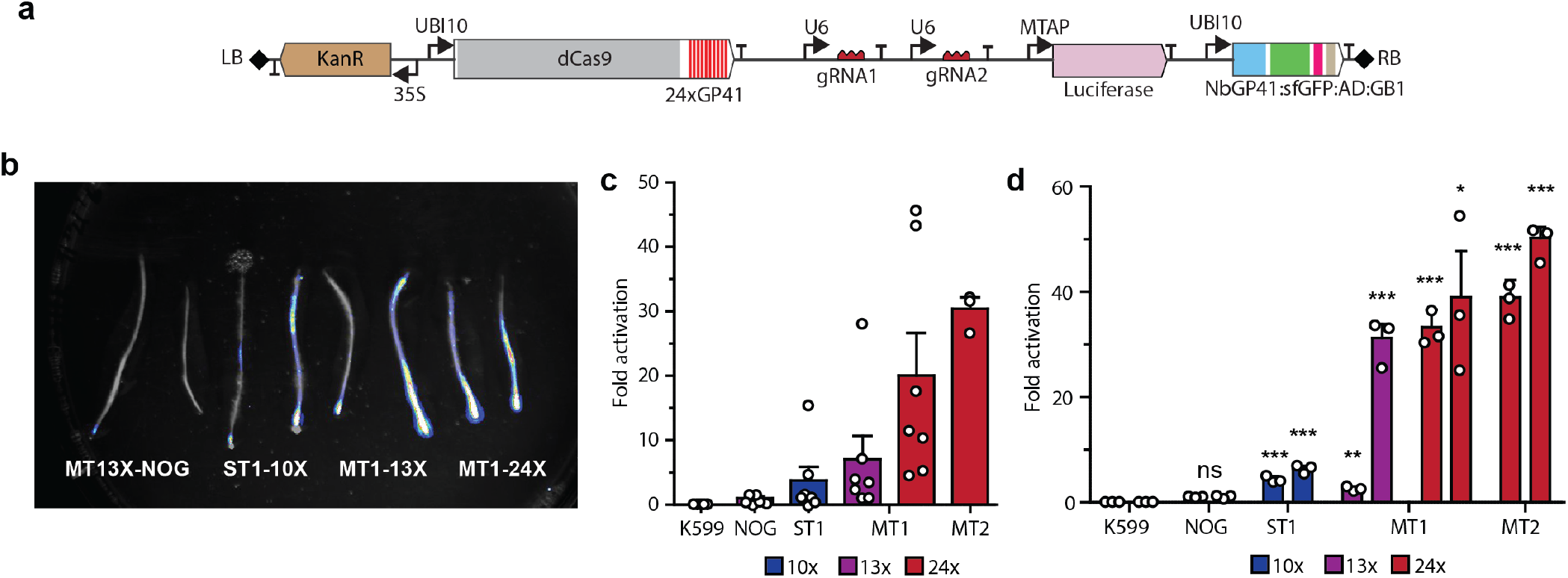
Activation of a luciferase reporter by MoonTag in tomato hairy roots. (a) T-DNA of the binary vector constructed to express MoonTag activating a luciferase reporter in tomato. The genes encoding the different components of MoonTag are indicated. dCas9-24XGP41 and NbGP41-sfGFP-VP64-GB1 are driven by the Arabidopsis UBI10 promoter. The gRNAs, indicated as gRNA1 and gRNA2, are driven by the Arabidopsis U6 promoter. Expression of the selectable marker NPTII is driven by the 2X 35S promoter. The Luciferase reporter is driven by synthetic promoter with binding site for gRNA1 and gRNA2. Right (RB) and left borders (LB) of the T-DNA are represented by grey rhombi. (b) Luminescence signal captured with a CCD camera superimposed on a bright field image of tomato hairy roots transformed with the indicated constructs. (c) Expression of Luciferase measured by qRT-PCR in the different hairy root lines one month after transformation. Each point in the graph represents an independent transformation event. (d) Expression of Luciferase after 10 months of subculture in two independent hairy root lines (left and right bars) transformed with the constructs indicated on the x-axis. Each point in the graph represents an independent RT-qPCR measurement. K599, tomato hairy root without any activator or Luciferase transgene. Gene expression was normalized to that of *SlACT2*.

### Activation of *FT* and *CLV3* by MoonTag-VP64 in *Arabidopsis thaliana*

In Arabidopsis, we generated transgenic plants expressing MoonTag and gRNAs that targeted the *CLAVATA3* (*CLV3*) and *FLOWERING LOCUS T* (*FT*) genes. For transformation with Agrobacterium, each construct contained the MoonTag components driven by the UBI10 constitutive promoter, two sgRNAs targeting the promoter or either *FT* or *CLV3*, and a selectable marker conferring kanamycin resistance. After transformation, kanamycin resistant seedlings expressing MoonTag targeting *FT* (16 lines) and *CLV3* (13 lines) were obtained. Expression of *FT* in the 16 lines ranged between 2 to 20 fold higher than that of the wild type (Supplementary Figure S8). Expression of *CLV3* in the 13 lines targeting *CLV3* was between 200 to 400 fold higher than in wild type (Supplementary Figure S8). Arabidopsis plants targeting *FT* or *CLV3* for overexpression where allowed to self and lines were further propagated to isolate homozygous lines. Two homozygous FT lines (FT10 and FT16) showed between 20 to 50 fold induction of *FT* compared to the wild type (Col-0) or the no-guide control (NOG)(Figure 3b). These lines have a clear early-flowering phenotype expected from prodigious *FT* expression (Figure 3b-c). Similarly, homozygous CLV3 lines showed a phenotype consistent with perturbation of the CLV3/WUS signaling pathway, including smaller size, loss of meristematic cells, and reduced leaf number (Figure 3f).

**Fig. 3.**
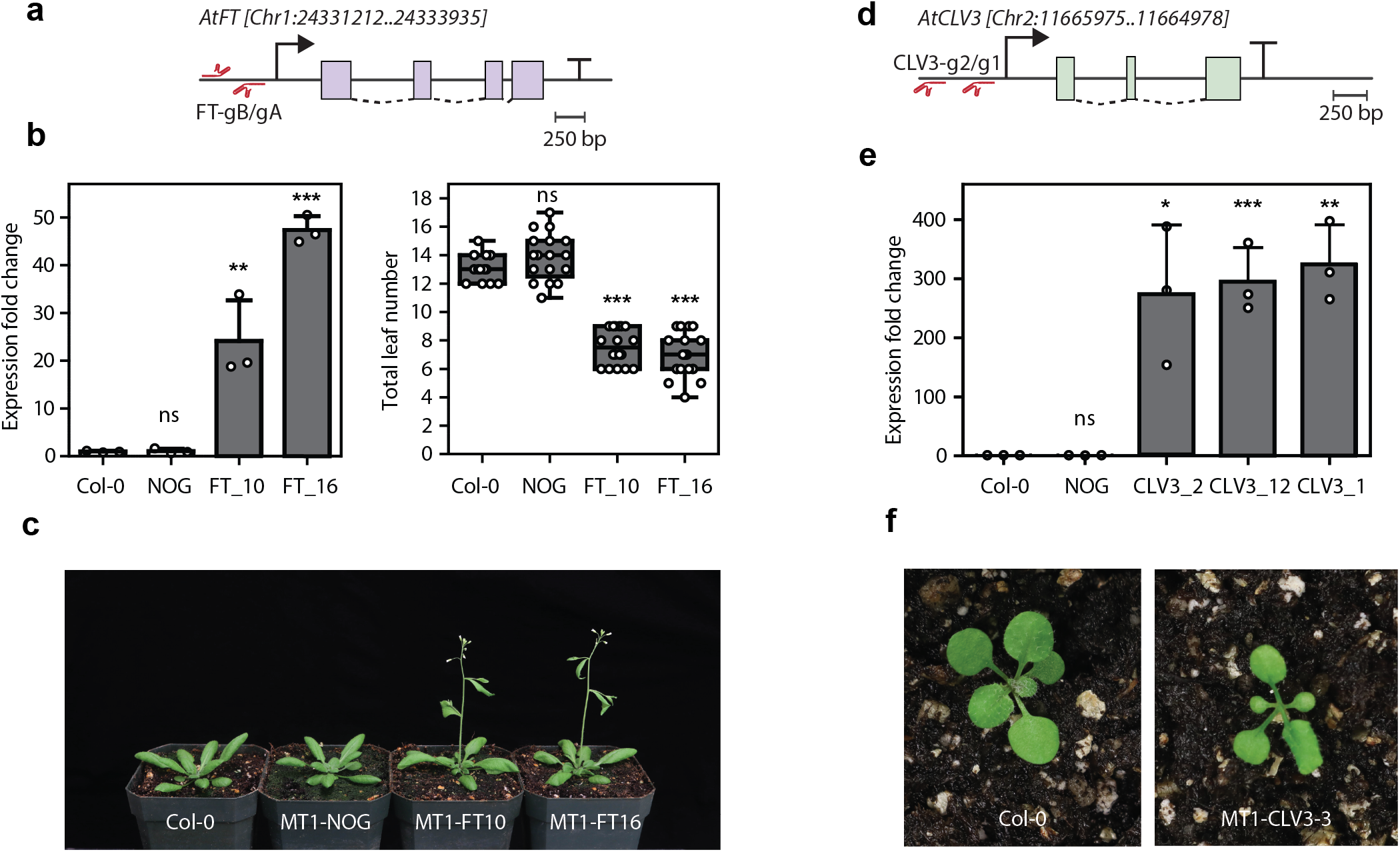
MoonTag is capable of activate endogenous genes in transgenic Arabidopsis plants. (a) Diagram of the *FT* gene showing the position of the gRNAs in relation to the transcription start site. (b) left: Expression of the FT gene in the indicated transgenic lines. right: Total rosette leaf number until flowering shown by each transgenic line. Expression of the FT gene was quantified in 6 day-old seedlings from homozygous lines collected at Zeitgeber time (ZT) 15. Transcript levels were normalized against *TUB2*. (c) Flowering phenotype of the indicated transgenic lines d) Diagram of the *CLV3* gene indicating the position of the gRNAs designed to activate this gene. e) Expression of *CLV3* in 7 day-old seedlings of the indicated lines. Expression of the *CLV3* gene was quantified in whole seedlings from Col-0 and plants homozygous for the transgenes. Transcript levels were normalized against *TUB2*. (f) Representative phenotype resulting from the overexpression of *CLV3*.

### Temperature effect in MoonTag-mediated gene activation

The endonuclease activity of catalytically-competent Cas9 is dependent on the temperature in which the target organism or cell-line is growing (21). While targeted mutagenesis via Cas9 is efficient at temperatures higher than 37 °C, it becomes gradually impaired at lower temperatures (21). The effect of temperature in the gene activation mediated by dCas9-based PTAs remains poorly studied. In order to investigate how temperature impacts MoonTag-mediated activation, we subjected either (i) hairy roots containing a MoonTag-activated luciferase reporter or (ii) Arabidopsis seedlings containing a MoonTag-activated endogenous gene (*CLV3*), to growth in different temperatures and measured the changes in activation relative to the growth at 24 °C. As shown in Figure 4, robust overexpression of target genes in both systems is observed from 4 °C to 28 °C. There is a general trend of decreased strength of overexpression at lower temperatures, despite the observation that expression of the MoonTag components remains relatively constant (Supplementary Figure S9). The boost in expression of luciferase in tomato hairy root cultures at 4 °C is explained by an increased expression of MoonTag components at this temperature, possibly the result of a cold-shock responsiveness of the UBI10 promoter (Figure 2a) (22). These observations show MoonTag to be a powerful activator of gene expression across a broad window of temperatures, albeit with a temperature-dependence.

**Fig. 4.**
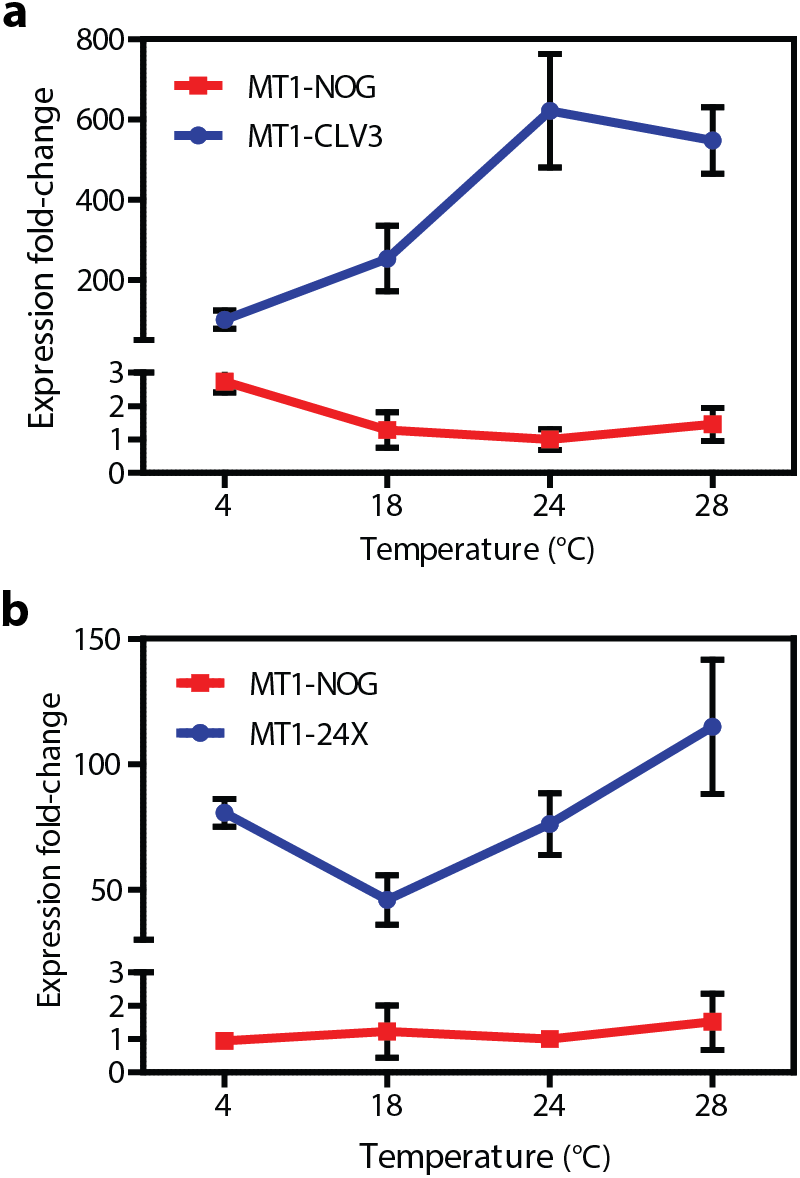
MoonTag activation is reduced but still effective at lower temperature in Arabidopsis and tomato. (a) Activation of *CLV3* by a complete MoonTag activator (blue) or no-sgRNA control (red) in Arabidopsis seedlings incubated at different temperatures. (b) Luciferase expression driven by a complete MoonTag activator (blue) or no-sgRNA control (red) in tomato hairy roots incubated at the indicated temperatures. Transcript levels were normalized to that of *TUB2* and *SlACT2* for Arabidopsis and tomato, respectively, and shown relative to the no-sgRNA control (NOG) grown at 24 °C. Values are the mean of three biological replicates ± standard deviation.

## Discussion

The diverse array of recently described CRISPR/Cas-based activators allow for targeted control of gene expression with minimal off-target effects in the transcriptome (1, 6, 8, 9). Multiple sgRNAs can be delivered to coordinate the overexpression, repression, or both, of multiple gene targets. Because this multi-gene targeting requires only the addition of relatively small sgRNA sequences to genetic constructs, CRISPR/Cas-based activators show promise in modulating multi-locus traits. Applications in plants include addressing basic research in plant biology, metabolic reprogramming, engineering beneficial traits in crops, or creating barriers to sexual reproduction and gene flow between engineered and wild type plants (15, 23–28). The MoonTag activators described here perform well in stable transgenic plants and are a suitable platform for developing such applications.

Genetic context effects can lead to unpredictable behavior of engineered genetic parts or devices when they are integrated to more complex systems. These can be further subcategorized into compositional context effects, host cell context effects, and environmental context effects (29). All CRISPR/Cas-based activators developed to date recruit multiple genetic components (DNA-binding domains, activation domains, modular interaction domains, etc.) from diverse organisms or viruses into an activator complex, which is typically used in a host cell that did not evolve to functionally express these components. While this can be mitigated in part by sourcing activator components from more closely-related lineages (18), this is not always possible.

We suspect that host-cell context effects pertaining to scFv expression underlies our observation that SunTag systems did not function well in stably transformed Setaria lines. Intracellular expression of scFv antibodies is challenging, mainly due to the difficult to form the disulfide bonds necessary for proper folding in the reducing environment of the cytoplasm leading to lost of antigen-binding, protein aggregation and degradation (30–32). Indeed, in the original SunTag system, even tough a cytoplasm-soluble form of scFv was utilized (33), it sill required the addition of solubility tags (sfGFP and GB1) to the scFv component in order to avoid aggregation in mammalian cells (8). Thus, difficulties to express the scFv component of SunTag in Setaria may likely originate from the lack of proper folding of the scFv component. Furthermore, the differences we observed here between expression of scFv with and without the sfGFP in transgenic versus protoplast transient systems may be the result of scFv remaining unfolded to different degrees in both systems. Low activation levels with SunTag have also been observed in other transient systems such as with *N. benthamiana* agroinfiltration and rice protoplasts (11, 15).

Recruitment of multiple activation domains also require the use of components/domains that are usually sourced from divergent organisms or evolved to function in other cellular compartments. This makes it difficult for some of the components to be properly expressed in plants. As we shown here, this seems to be the case of the SunTag version we tried to transform into Setaria. The scFv component of SunTag we used, scFv-VP64-GB1, appears to be toxic when expressed in Setaria transgenic plants, even tough it worked well when transfected into protoplasts. In contrast a SunTag version using sfGFP, scFv-sfGFP-VP64, was demonstrated to be readily expressed in *A. thaliana* and to lead to high levels of gene activation (14), even tough in our protoplast transient system in Setaria lead to only low activation levels.

To avoid the issues inherent to the expression of scFv antibodies in plants but to maintain the efficient AD recruitment system of SunTag, we replaced the scFv antibody by a nanobody. We adapted the NbGP41 nanobody-GP41 peptide pair from the single molecule imaging reported by Boersma and colleagues (16) to create the MoonTag activator. Nanobodies are smaller than scFv fragments, naturally monomeric, contain a single disulfide bond that may not be necessary for proper folding and activity, thus making them likely to express better in eukaryotic cells (34, 35). This seems to be the case for the VP64 fusion with llama nanobody NbGP41 (NbGP41-sfGFP-VP64-GB1) which we were able to efficiently express in Setaria protoplasts and in transgenic plants of Setaria, tomato and Arabidopsis. In Setaria and tomato RNA expression levels of NbGP41-sfGFP-VP64-GB1 are usually higher that the highly expressed reference genes. Bright fluorescence from the sfGFP component could be also observed in the nuclei of transgenic plants. Additionally, expression of NbGP41-sfGFP-VP64-GB1 is higher than that of scFv-sfGFP-VP64 in few tomato hairy roots lines expressing MoonTag with gRNAs (MT1-13X and MT1-24X), and even when comparing two lines with the same expression levels of the antibody fusion, NbGP41 show brighter GFP fluorescence than scFv. Higher expression and GFP brightness of the AD fused to the nanobody GP41 suggest that this nanobody may fold better, and therefore, to accumulate to higher levels without the deleterious effects coming from expressing improperly folded proteins in the cytoplasm.

Besides improved expression of the nanobody-VP64 fusion component, making it to be less likely to cause toxicity when expressed in plant cells, we demonstrate that MoonTag is a powerful activator in the monocot plant Setaria and the eudicot plants, Arabidopsis and tomato. MoonTag-mediated activation in Setaria protoplasts is similar to that of the SunTag system and and can surpass it by using 24 copies of the GP41 peptide. A similar tendency was observed in tomato hairy roots where activation of a Luciferase reporter was achieved to higher levels with 13X and 24X copies of GP41 in MoonTag compared to SunTag expressed with only 10X copies of GCN4. In Arabidopsis, where a MoonTag version with 24 copies of the GP41 peptide was used, we observed higher levels of overexpression of the *CLV3* target (~250 fold) than those observed by Papikian an colleagues using SunTag with 10 copies of the GCN4 peptide (~ 100-150 fold) (14). In a similar way, activation of *FT* by MoonTag reached higher levels (~50 fold) compared to activation of the same gene using the SAM activator with MS2-VP64 (~35 fold) (13). Only the recently described TREE system-like activator, ACT3.0, seems capable of higher activation levels of the *FT* gene (130-fold to 240-fold) (15). However, as we recently shown activation of MoonTag, and other PTAS, can be increased with the use of novel ADs from plants origin (18). Furthermore, just as with SunTag, the NbGP41 nanobody-GP41 peptide pair, could be used to replace the scFv antibody-GCN4 peptide pair, in ACT3.0. Thus, MoonTag activation levels, par to second-generation activators, could be further improved by the use of more active ADs or by adapting MoonTag to novel PTA architectures that will benefit from the suitability of its components for expression in plant cells.

Coming from *Streptococcus pyogenes*, an organism with an optimal growth temperature of 40 °C (36), the nuclease activity of CRISPR Cas9 proteins may be partly impaired in organisms growing at temperatures lower than 37 °C such as plants (21). This temperature effect may be circumvented by limited high temperature treatments that lead to improved mutagenesis efficiency of Cas9 in plants (37–40). CRISPR activation, in contrast, rather depends on the binding of dCas9 to its target region but not its nuclease activity.

In addition, sustained activation requires the dCas9 protein to be constantly bound to its target region. The temperature requirements for binding must be different than those for cutting DNA, since binding does not depend on the nuclease activity. Indeed, at the normal growing temperature (~22-24 °C) MoonTag, and other CRISPR activators, seem to been able to efficiently activate their target genes suggesting that dCas9 can readily bind to their target loci. At lower temperatures, however, activation mediated by MoonTag is significantly reduced in Arabidopsis transgenic plants and tomato hairy roots. In our conditions this temperature impairment of gene activation took only 24 hours to manifest, suggesting that in field conditions, temperatures decreasing at night or few days of lower temperatures could have a marked effect in gene activation. We did not observed any changes in expression of MoonTag components or the gRNA that could account for our temperature observations suggesting that either DNA binding, AD recruitment or AD activation itself may be the cause of reduced activation at low temperatures. The Cas9 ribonucleoprotein is capable of binding DNA at temperatures as low as 4 °C in vitro (41), however, whether the DNA binding of activity dCas9 is similar to that of Cas9 or if it behaves similarly in vivo, as in vitro, remains to be determined. Currently we do not know how lower temperature affects other types CRISPR activators, but given the similarities between MoonTag and SunTag it is likely that a similar trend may happen with the later activator.

## Materials and methods

### Plant material

*Setaria viridis* accession ME034 was used in this study. Setaria plants were grown in growth chambers under a 12:12 h light/dark cycle, 31 °C/22 °C day/night temperatures with a light intensity of 400 PAR. When harvested seeds were younger than 6 months-old they were treated with gibberellic acid to break dormancy and promote germination as described before (42). *Arabidopsis thaliana* ecotype Col-0 was used in all experiments. Arabidopsis was grown at under a 16:8 h light/dark cycle, 22 °C/20 °C day/night temperature with a light intensity of 120-150 μmol/m^2^. When germinated on plates seedlings were surface-sterilized with 20% bleach for 10 min and planted on 1/2MS20 media (half-strength Murashige and Skoog basal salt mixture, 20 g/L sucrose, pH 5.8). *Solanum lycopersicum* cultivar M82 was used to generate hairy roots. Tomato seeds were sterilized in 50% bleach for 15 minutes and then germinated in jars containing MS30 media (full-strength Murashige and Skoog basal salt mixture, 30 g/L sucrose, 0.5 g/L 2-(N-morpholino)ethanesulfonic acid, 10 g/L agar, pH 5.8). All in vitro culture was carried out at 24 °C under a 16:8 h light/dark cycle with a light intensity of 150 μmol/m^2^.

### vector construction

The dead Cas9 coding sequence (dCas9) including nuclear localization signals was obtained from pEG302 22aa SunTag VP64 nog (Addgene plasmid #120251). DNA fragments encoding different copy numbers of the GP41 peptide separated by a 5 amino acids GS linkers were derived from the plasmid 24×MoonTag-kif18b-24×PP7 (addgene plasmid #128604) and used to make a translational fusion downstream to the dCas9 sequence. A DNA fragment encoding the GP41 nanobody was cloned from the plasmid Nb-gp41-GFP (MoonTag-Nb-GFP) (addgene plasmid #128602) and used to replace scFv on the CDS of scFv-sfGFP-VP64-GB1 cloned from the plasmid pEG302 22aa SunTag VP64 nog. The CDS corresponding to dCas9-10XGCN4 and scFv-sfGFP-VP64-GB1 were obtained from pEG302 22aa SunTag VP64 nog. SunTag and MoonTag dCas9-peptide and antibody-AD fusions were assembled into Level 0 destination vectors using BsaI with the Phytobrick (J6-J9) overhangs. Level 1 constructs were assembled using Golden Gate with the BsaI restriction enzyme to generate transcriptional units driven by the promoters and terminators used in each construct (43). Guide RNAs were designed and expressed from U6 promoters, the *Oryza sativa* U6 (OsU6) promoter for Setaria and *A. thaliana* U6 (AtU6) promoter for tomato and Arabidopsis. The gRNA constructs were made using Golden Gate cloning as described before (44). The backbone for the gRNA driven by the OsU6 promoter was pMOD-B2520 and the backbone for the gRNA driven by AtU6 was pMOD-B2515. Level 1 constructs together with gRNA constructs were in turn assembled into binary vectors suitable for Monocot or dicot transformation (44).The list of gRNAs sequences used in this work is listed in Supplementary Table S1.

### Protoplast isolation and transfection

Protoplasts from Setaria leaves were isolated as described before (45). Transfection was carried out using polyethylene glycol (PEG). For transfection of protoplasts for RNA analysis, 500000 cells were mixed with plasmid DNA corresponding to the different constructs (10 μg per construct) in 20% PEG for 10 minutes. After transfection, protoplasts were incubated at room temperature in the dark for 16 to 18 hours. Protoplast transfection for luciferase assays was carried out in 96-well culture plates using 100000 cells per well and 2 μg of plasmid DNA for each construct. A plasmid expressing Renilla luciferase from a 35S promoter was added to be used in downstream analysis to normalize the activity of firefly luciferase (18).

### Luciferase assay of protoplasts

Luciferase assay were performed as described previously (45)(18). Protoplasts were collected by centrifugation, resuspended in 20 μl of passive lysis buffer (Promega) and lysis allow to happen for 15 minutes at room temperature with shaking at 40 rpm. Firefly and Renilla Luciferase activities in the lysate were then determined with a Dual-Luciferase Reporter Assay System (Promega) and a GloMax explorer plate reader (Promega) following the manufacturer instructions. Firefly luciferase activity in the different treatments was normalized to that of Renilla Luciferase.

### RNA isolation

RNA was isolated from different plant tissues using the Trizol reagent (Thermo Fischer Scientific) following the manufacturer instructions. For Setaria plants and tomato hairy roots, 50-100 mg of tissue were extracted with 1 mL of Trizol whereas for Setaria protoplasts, 750 μL of Trizol was used for 500000 cells. To analyze Arabidopsis *FT* expression, RNA was isolated from 4-5 pooled six-day-old seedlings grown under long-day (LD) conditions and harvested at Zeitgeber time (ZT) 15. RNA for Arabidopsis *CLV3* RT-qPCR analysis was isolated from 7 day-old seedlings grown in LDs. RNA was resuspended in nuclease-free water (TAKARA) and then treated with DNAse using the TURBO DNA-free Kit (Invitrogen).

### Quantitative reverse transcription PCR (RT-qPCR)

RT-qPCR was carried out from the isolated RNA using the Luna Universal One-Step RT-qPCR Kit (New England Biolabs) following the manufacturer instructions. RT-qPCR for RNA samples from Setaria leaves, Arabidopsis seedlings and tomato hairy roots was done using 100-150 ng of RNA per reaction. For protoplasts, each reaction was performed using 25-35 ng of RNA. Expression of the genes tested in Setaria, tomato hairy roots and Arabidopsis were normalized to that of *SvGRAS, ACTIN2* (*SlACT2*), and *TUBULIN2* (*TUB2*), respectively. The primers used in all Rt-qPCR analysis are listed in Supplementary Table S2.

### Agrobacterium mediated plant transformation

Arabidopsis plants ecotype Col-0 was transformed with *Agrobacterium tumefaciens* GV3101 carrying the binary vectors of interest and using the “floral dip” method (46). After transformation seeds were harvested and putative transformants were identified by kanamycin selection in 1/2MS20 media containing 50 mg/L of kanamycin. Seedlings resistant to kanamycin were transferred to soil and grown until the plants set seeds.

#### Setaria viridis

ME034 cultivar was transformed using *Agrobacterium tumefaciens* strain AGL1 following the procedure described by Weiss and colleagues (42). After regeneration putative transgenic plants were transferred to soil and allowed to set seeds. Harvested seeds were then germinated in the presence of hygromycin (50 mg/L) and only resistance seedlings were transferred to soil for further analysis.

### Hairy root transformation

*Agrobacterium rizhogenes* K599 was used to induce hairy roots in the tomato cultivar M82. Transformation was carried out as previously reported (47). After transformation, hairy roots for all activators were maintained in MS30 media supplemented with 50 mg/L of kanamycin and 200 mg/L of timentin. Hairy roots produced by Agrobacterium K599 without any vector were maintained in MS30 supplemented with 200 mg/L of timentin.

### Temperature treatments

Homozygous seeds from a line expressing MoonTag with two gRNAs targeting *CLV3* (MT1-CLV3) and a line expressing MoonTag without gRNAs (MT1-NOG) were surface sterilized and placed in 1/2MS20 media supplemented with 50 mg/L of Kanamycin. Plates were incubated for 48 hours at 4 °C and then placed at 24 °C under a 16:8 h light/dark cycle with a light intensity of 150 μmol/m^2^. Seeds were allowed to germinate and grow for 6 days. Plates containing seeds were then placed into incubators a 4 °C, 18 °C, 24 °C and 28 °C in the dark for 24 hours after which samples were collected for RNA isolation. 4-5 seedlings were pooled into a single tube, flash-frozen in liquid nitrogen and stored at −80 °C until RNA isolation. For hairy roots, a line expressing MoonTag with two gRNAs targeting a synthetic promoter driving luciferase reporter (MT1-24X) and a line expressing MoonTag, the luciferase reporter but with any gRNAs (MT1-NOG) were used. a single hairy root was transferred to a single plate of MS30 media supplemented with 50 mg/L of kanamycin and allowed to grow for seven days at 24 °C. Plates were then transferred to incubators set to the temperatures indicated above. 24 hours later the roots were placed in tubes, flash-frozen in liquid nitrogen and stored at −80 °C until RNA isolation.

## ACKNOWLEDGEMENTS

MHZ, JACM, and MJS are supported by the USDA grant 2018-33522-28747 and are supported by the Advanced Plant Technologies program, DARPA Award HR001118C0146. MHZ is supported by an NIH NIGMS Biotechnology Training grant NIHT32GM008347.

## AUTHOR CONTRIBUTIONS

JACM, MZ and AS designed experiments and collected data. JACM, MZ, and MJS analyzed the data. JACM and MJS wrote the manuscript.

## Supplementary Note 1: Supplementary Figure 1

**Fig. S1. Supplementary Figure 1.**
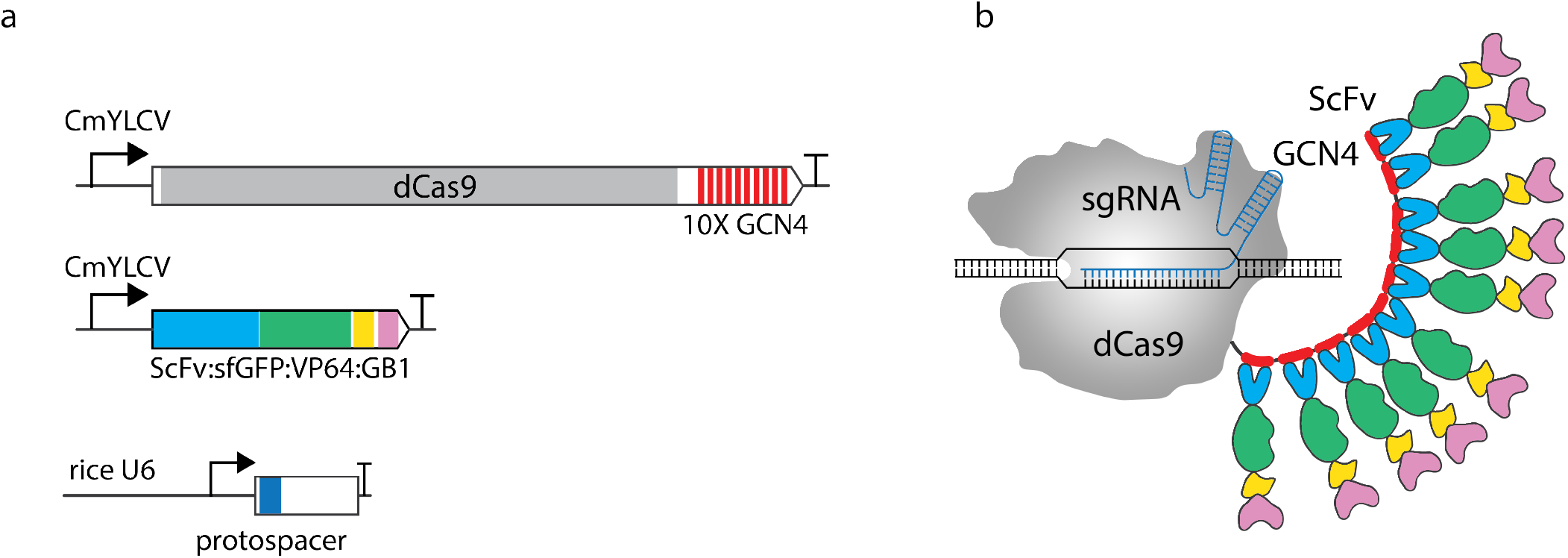
Schematic representation of the SunTag activator. (a) Schematic diagrams of the expression constructs encoding the SunTag components. The DNA binding component consist of dCas9 fused to 10 copies of the GCN4 peptide (dCas9-10XGCN4); an activation module comprised of the scFv antibody fused to sfGFP (super folder GFP), the VP64 activation domain and the GB1 solubility tag (NbGP41:sfGFP:VP64:GB1); and the sgRNA expression cassette driven by a U6 promoter. (b) When expressed in plant cells dCas9-10XGCN4 is able to bind to DNA guided by the sgRNA. The GCN4 peptides in dCas9-10XGCN4 are bound by the scFv antibody of scFv-sfGFP-VP64-GB1 recruiting up to 10 copies of the VP64 activation domains.

## Supplementary Note 2: Supplementary Figure 2

**Fig. S2. Supplementary Figure 2.**
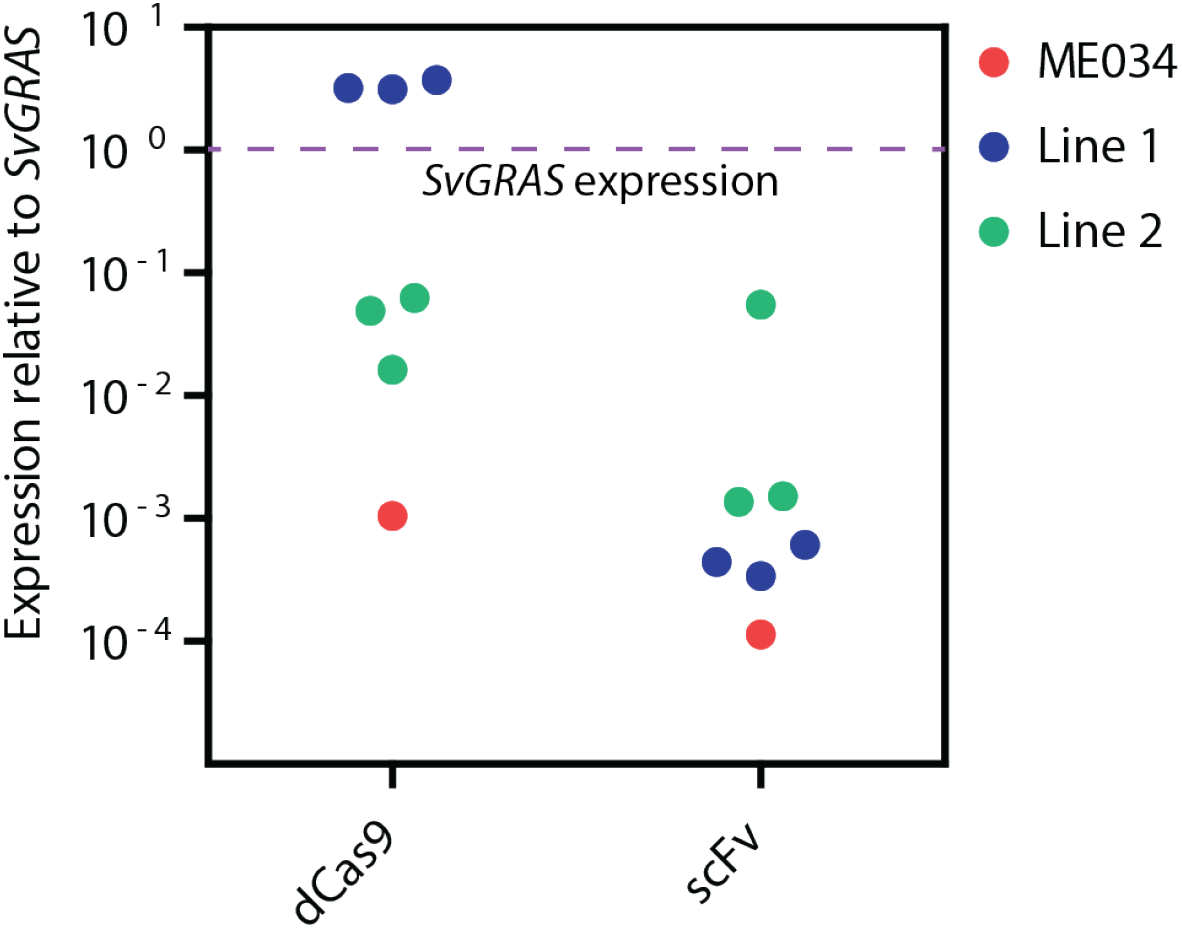
Expression of SunTag components in Setaria. Expression of dCas9-24XGCN4 (dCas9) and scFv-VP64-GB1 (scFv) components of SunTag was determined in two transgenic lines. RNA was isolated from leaves of three homozygous plants from each line. Expression of each component is shown relative to that of *SvGRAS*.

## Supplementary Note 3: Supplementary Figure 3

**Fig. S3. Supplementary Figure 3.**
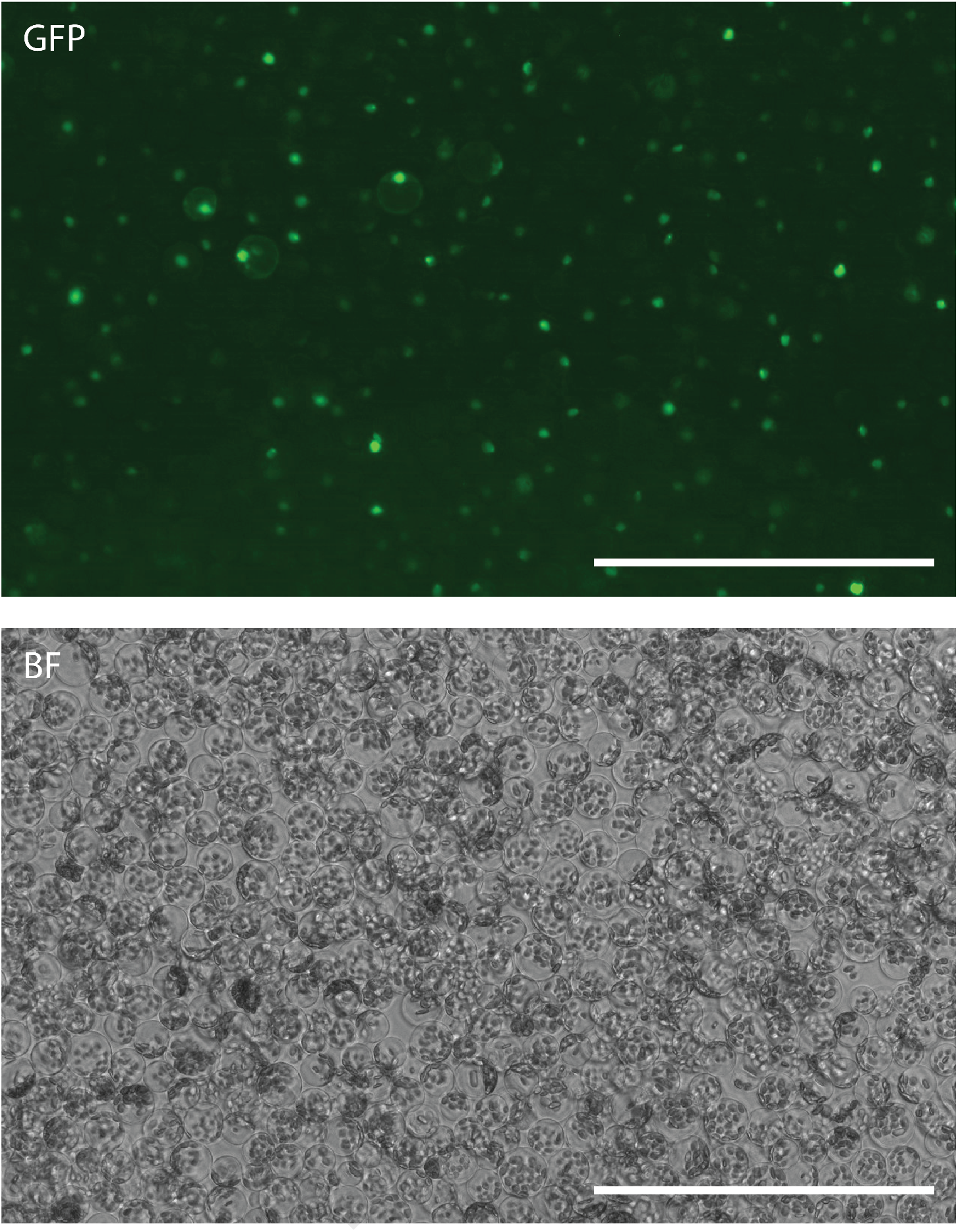
Expression of the nanobody NbGP41 fusion with sfGFP in Setaria protoplasts. The construct NbGP41-sfGFP-VP64-GB1 under the control of the cmYLCV promoter was transfected into Setaria protoplasts. GFP expression was analyzed 24 hours after transfection using fluorescent (GFP) and bright field (BF) microscopy. bar size 200 μm.

## Supplementary Note 4: Supplementary Figure 4

**Fig. S4. Supplementary Figure 4.**
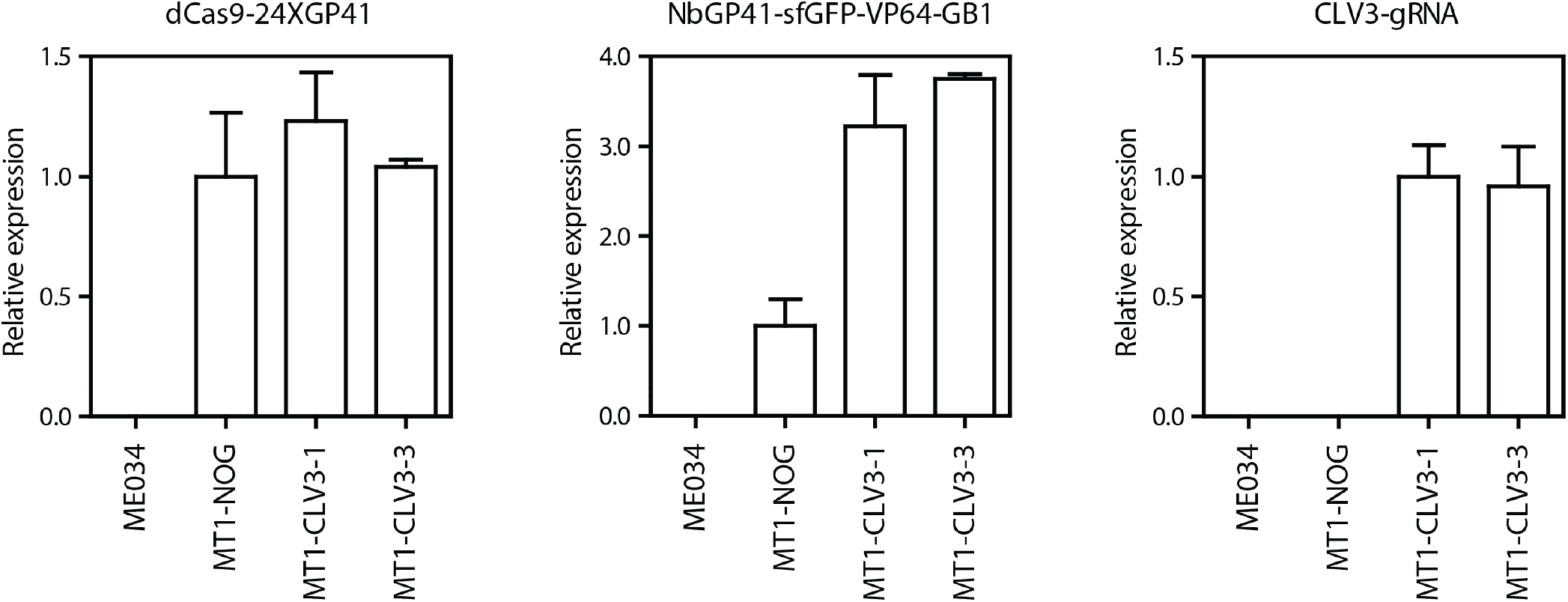
Expression of MoonTag components in the wild type ME034, the no-sgRNA control (MT1-NOG) and two homozygous Setaria transgenic plants. Left panel: Expression of dCas9-24XGP41 in two homozygous T2 Setaria transgenic plants. Middle panel: Expression of NbGP41-sfGFP-VP64-GB1 in homozygous T2 Setaria transgenic plants. Right panel: Expression of the gRNA3 targeting *SvCLV3* in homozygous T2 Setaria transgenic plants. For ME034, MT1-NOG and each transgenic line, MT1-NOG, MT1-CLV3-1 and MT1-CLV3-3, RNA from the leaves of 3 homozygous individuals were analyzed. Expression was normalized to *SvGRAS* and then relative to MT1-NOG in (a) and (b) and to MT1-CLV3-1 in (c).

## Supplementary Note 5: Supplementary Figure 5

**Fig. S5. Supplementary Figure 5.**
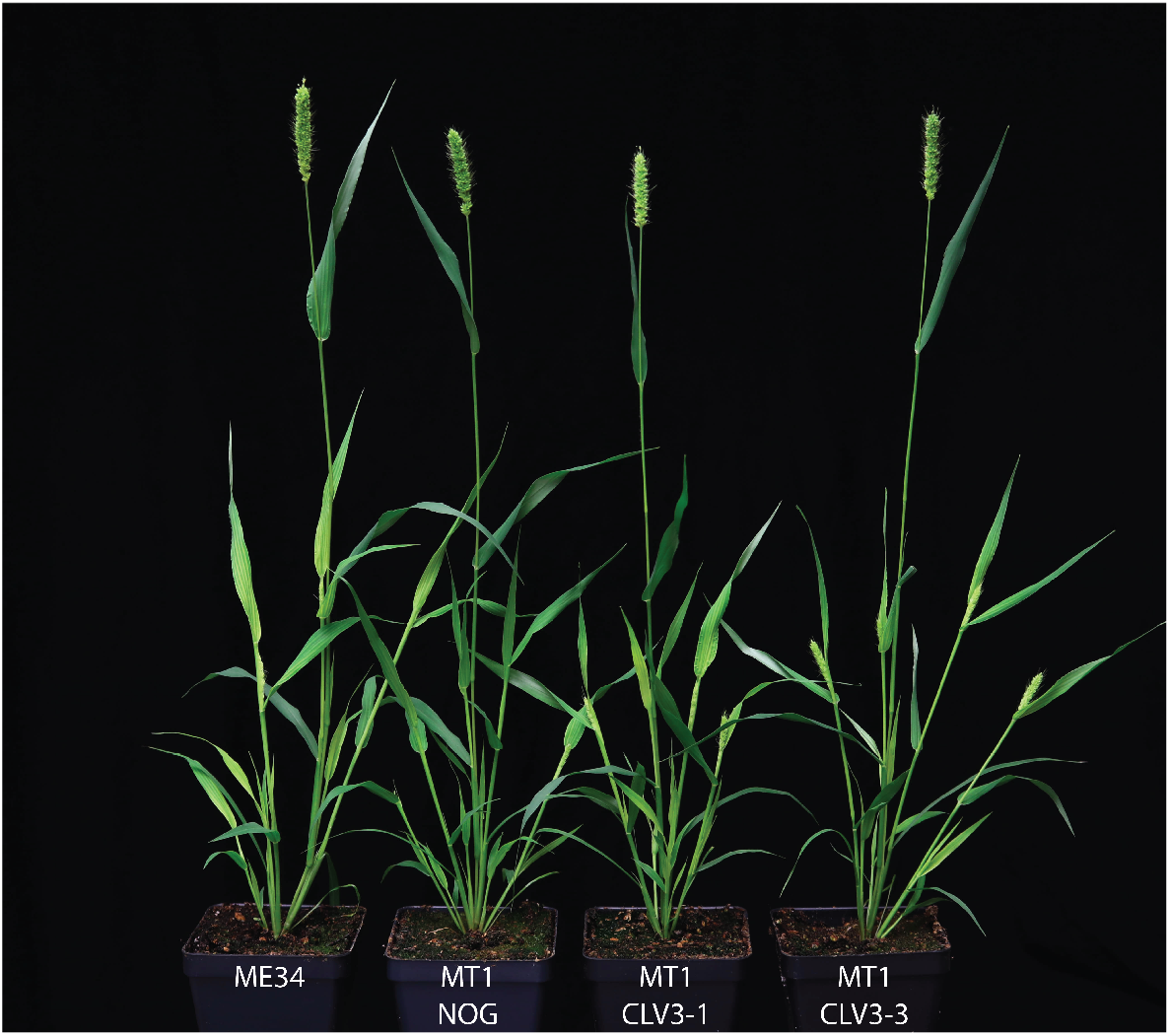
Phenotype of Setaria wild type ME034, the no-sgRNA control (MT1-NOG) and two homozygous Setaria transgenic plants expressing MoonTag with the gRNA3 targeting *SvCLV3*. Plants were grown in a growth chamber under a 12:12 h light/dark cycle, 31 °C/22 °C day/night temperatures with a light intensity of 400 PAR.

## Supplementary Note 6: Supplementary Figure 6

**Fig. S6. Supplementary Figure 6.**
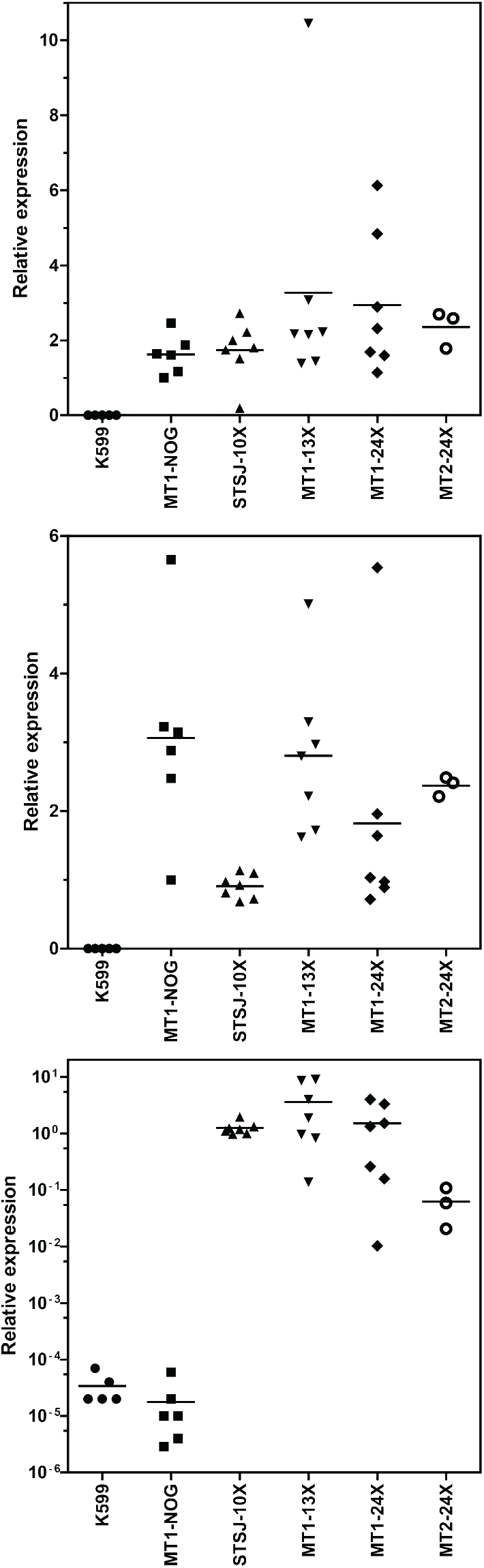
Expression of dCas9-peptide fusion, antibody fusion and gRNAs in the indicated lines of tomato hairy roots. Top panel: Expression of antibody fusions. NbGP41-sfGFP-VP64-GB1 for MT1-NOG, MT1-13X and MT1-24X; NbGP41-VP64-GB1 for MT2-24X, and scFv-sfGFP-VP64-GB1 for STSJ-10X. Middle panel: Expression of the dCas9-peptide fusions. dCas9-13XGP41 for MT1-NOG and MT1-13X; dCas9-24XGP41 for MT1-24X and MT2-24X; and dCas9-10XGCN4 for STSJ-10X. Lower panel: Expression of the gRNAs targeting the synthetic promoter MTAP1. K599, tomato hairy root without any activator or luciferase transgene; MT1-NOG, no-sgRNA control with dCas9-13XGP41, NbGP41-sfGFP-VP64-GB1 and the MTAP1 promoter driving a luciferase reporter. The scale in the lower panel is set as logarithmic due to the wide range of expression levels of the gRNAs. Expression is normalized to *SlACT2* and shown relative to MT1-NOG for the top and middle panels and to STSJ-10X for the lower panel. Each data point is an independent hairy root line. The average of the values in each column is indicated by a solid line.

## Supplementary Note 7: Supplementary Figure 7

**Fig. S7. Supplementary Figure 7.**
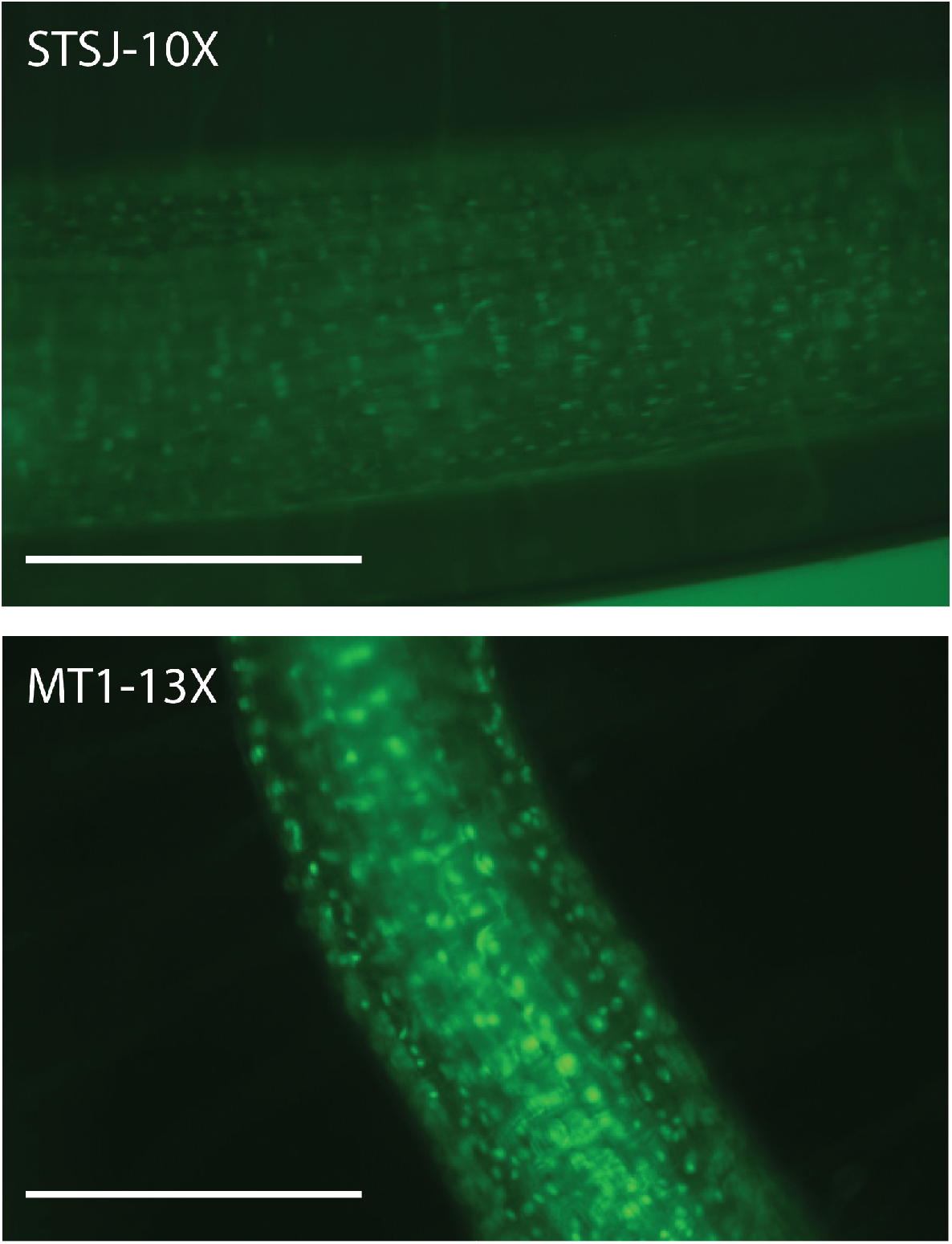
sfGFP fluorescence of the antibody fusion NbGP41-sfGFP-VP64-GB1 of MoonTag (MT1-13X) and scFv-sfGFP-VP64-GB1 of SunTag (ST1-10X) in tomato hairy roots. Pictures were taken from approximately the same length of the root relative to the root tip, using the same UV light illumination. bar size 400 μm

## Supplementary Note 8: Supplementary Figure 8

**Fig. S8. Supplementary Figure 8.**
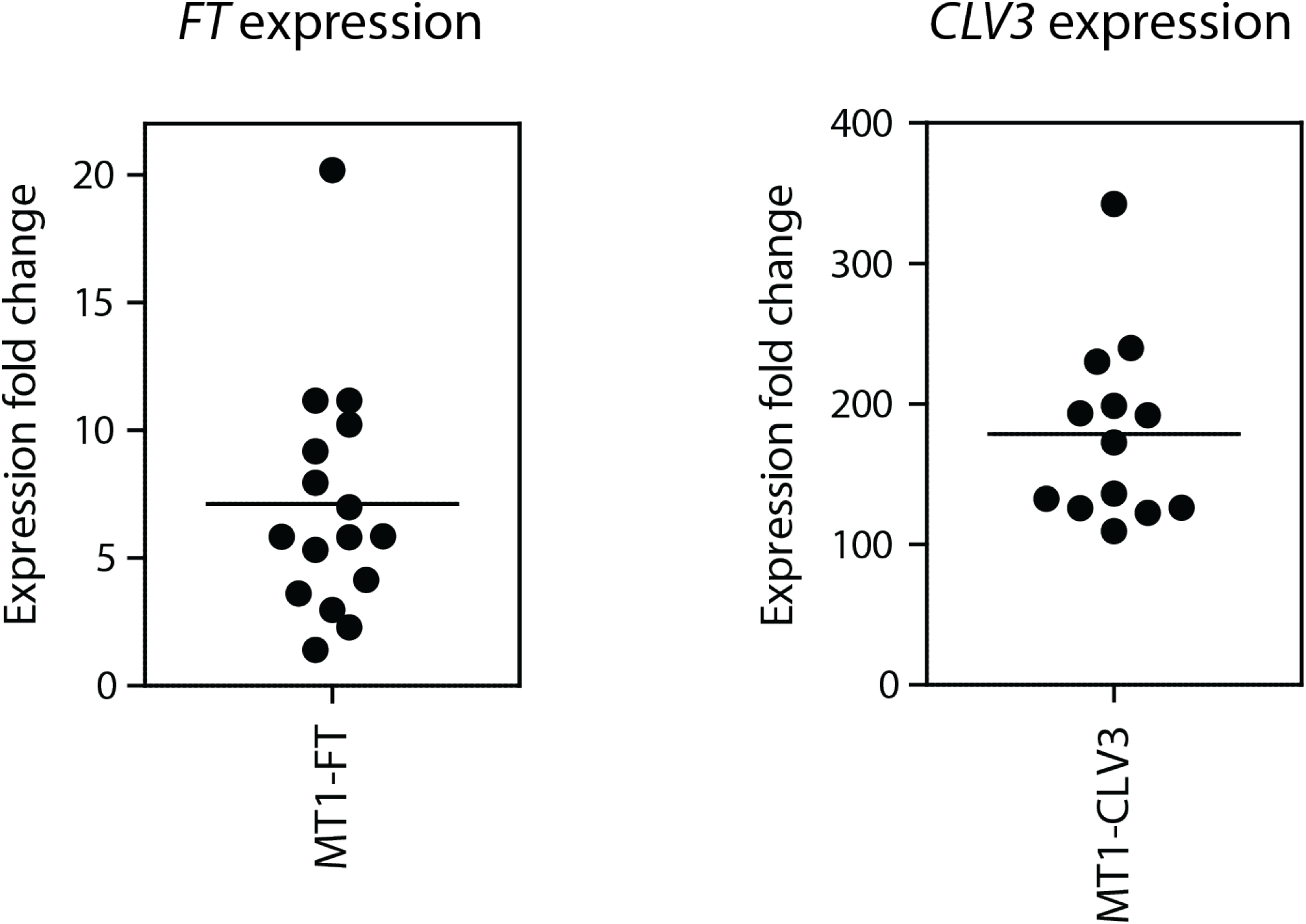
Expression of the indicated genes in T1 Arabidopsis transgenic plants. Each dot corresponds to an independent transgenic line. Expression of the *FT* gene was quantified in 6 day-old seedlings from homozygous lines collected at Zeitgeber time (ZT) 15. Expression of *CLV3* was quantified in 7 day-old seedlings. RNA was isolated from 4-5 pooled seedlings. Expression is normalized to that *TUB2* and shown relative to Col-0. Average fold-change expression in all lines is shown with a solid line.

## Supplementary Note 9: Supplementary Figure 9

**Fig. S9. Supplementary Figure 9.**
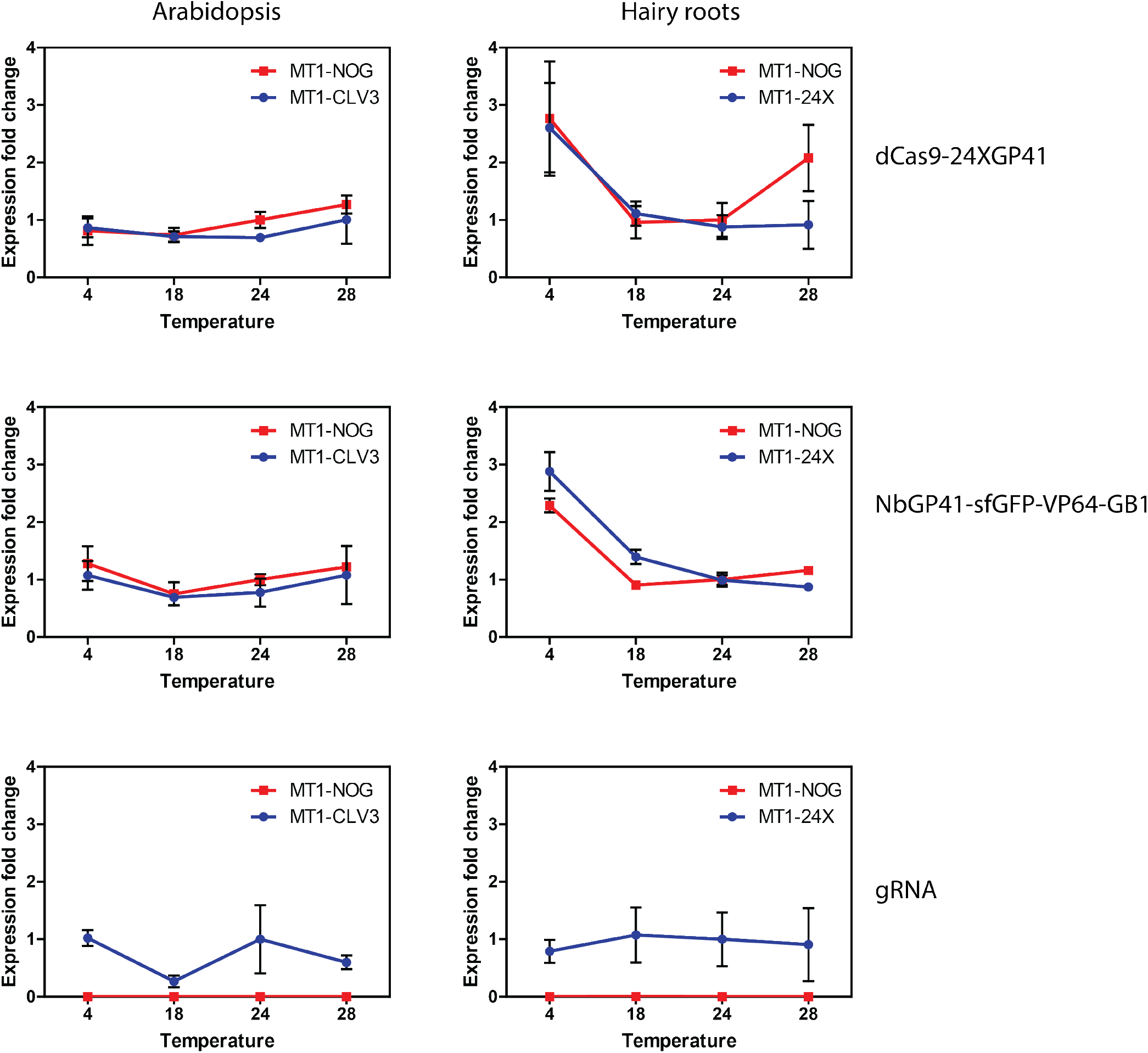
Expression of MoonTag components and gRNAs at the indicated temperatures in Arabidopsis and tomato hairy roots. Arabidopsis MT1-CLV3 line (blue) expresses the MoonTag activator with two gRNAs targeting the *CLV3* gene. Tomato hairy root line MT1-24X (blue) expresses MoonTag with two gRNAs targeting the MTAP1 promoter driving the luciferase reporter. MT1-NOG (red) is a no-sgRNA control in Arabidopsis or tomato hairy roots. Expression was normalized to that of *TUB2* for Arabidopsis and *SlACT2* for tomato hairy roots, and shown relative to the no-sgRNA control (MT1-NOG) grown at 24 °C. Values are the mean of three biological replicates ± standard deviation. Temperature treatments were carried out as indicated in methods.

## Supplementary Note 10: Supplementary Table 1

**Table S1. Supplementary Table 1.**
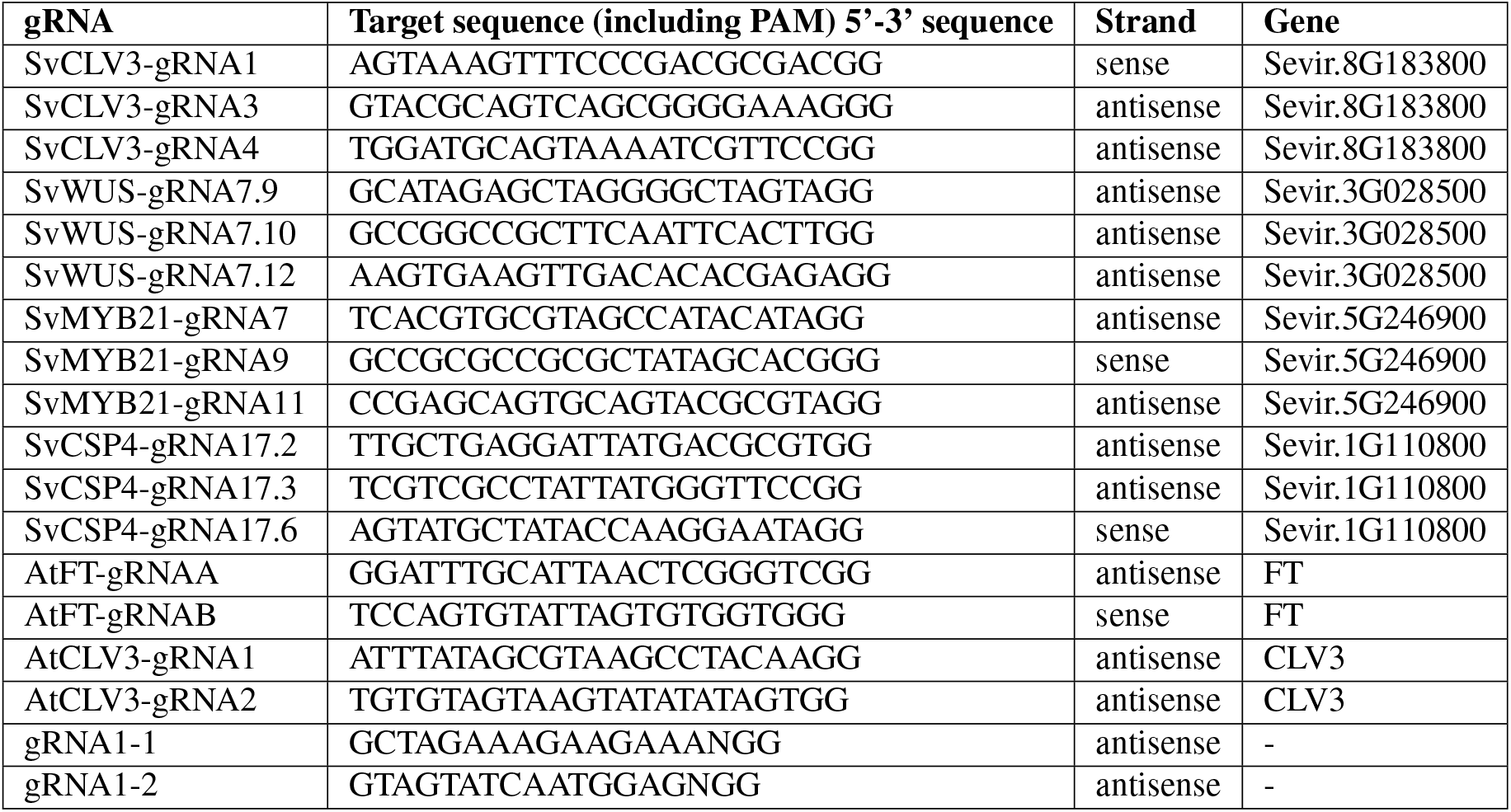
Sequences of the gRNAs used in this this study.

## Supplementary Note 11: Supplementary Table 2

**Table S2. Supplementary Table 2.**
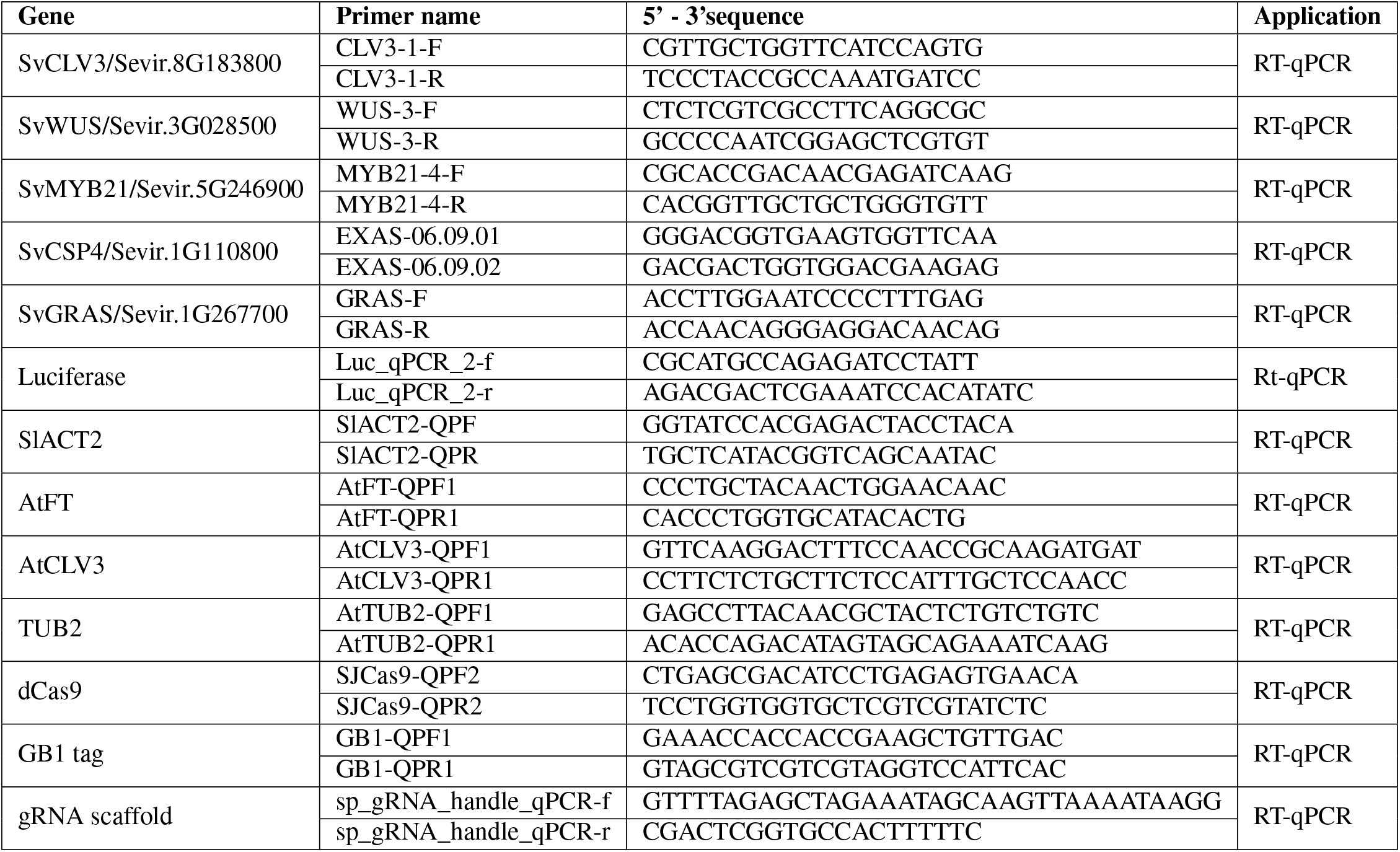
List of primers used in this this study.

